# Using X-ray microtomography to uncover the visual world of ancient and historical insects

**DOI:** 10.1101/290841

**Authors:** Gavin J. Taylor, Stephen A. Hall, Johan A. Gren, Emily Baird

## Abstract

Animal eyes typically have specialized regions of high resolution and/or sensitivity that enable them to execute behavior effectively within their specific visual habitat. These specializations, and evolutionary changes to them, can be crucial for understanding an animal’s ecology. Techniques for analyzing visual specializations typically require fresh samples and are thus limited to studies on extant species. However, the cornea of invertebrate compound eyes is readily preserved, even in fossils. To compare and quantify vision in specimens from different time periods and habitats, and with different preservation states – fossilized in amber, dried or stored in alcohol – we developed a novel technique that uses X-ray microtomography to create high resolution 3D models of compound eyes from which their sensitivity, spatial resolution, and field of view can be estimated using computational geometry. We apply our technique to understanding how the visual systems of some of the smallest flying insects, fungus gnats, have adapted to different types of forest habitat over evolutionary time (~30 mya to today). Our results demonstrate how such investigations can provide critical insights into the evolution of visual specializations and the sensory ecology of animals in general.

## Introduction

Vision provides essential information for guiding the behaviour of many animals and is often critical for orchestrating both movement through, and interactions with, their environment (Land & Nilsson 2012). To meet the sensory demands of visually-guided behaviour, animal eyes typically contain specialized regions of high resolution and/or sensitivity that are optimized for acquiring information to guide specific behavioural tasks, such as pursuing conspecifics or visually identifying flowers. Another, less well-understood factor affecting visual adaptations is the environment itself because specific features, such as the illumination and presence of a clear horizon may vary dramatically – consider, for example, the difference between a bright open meadow and a dim cluttered rainforest (Cronin et al. 2014). To acquire the visual information necessary for guiding behaviour in these different environments, animal eyes are also likely to have habitat-specific visual specializations (e.g. Scales & Butler 2016). Understanding these visual adaptations, and evolutionary changes to them, is crucial for understanding an animal’s sensory ecology and its ability to adapt to both long- and short-term habitat changes.

The corneal structure of the arthropod compound eye is commonly used to identify visual specializations from the external size and distribution of the corneal facets (Ribi et al. 1989, Bergman & Rutowski 2016), and these are often preserved naturally on samples in taxonomic collections (Suarez & Tsutsui 2004). The diverse range of arthropod specimens existing in museum collections is thus an ideal resource for the comparisons needed to study whether and how eyes evolve habitat-specific adaptations, or other questions regarding the factors driving eye evolution. However, current methods for analysing the visual properties of corneas are unsuitable for such investigations because i) some methods require fresh tissue and are destructive, so they can only be applied to readily available and extant species (Schwarz et al. 2011), ii) most methods only provide a 2D representation of the compound eye and thus limit our ability to understand how it viewed its 3D world (Ribi et al. 1989, Bergman & Rutowski 2016). We recently developed a method to quantify and recreate the visual world of arthropod eyes using X-ray microtomography (microCT) – a popular tool for morphological and structural analysis in many areas of biology (Baird & Taylor 2017) – and applied it to extant insect specimens that had been specially prepared and stained (Taylor et al. 2019). Here, we present a further development to this method, which enable us to analyse the corneal morphology of naturally preserved specimens that are fossilized in amber or dried, as well as specimens preserved in alcohol. From this, we are able to quantify the vision of insects from taxonomic collections and in different states of preservation. Additionally, we develop a novel method for simulating the visual world of insects from the calculated visual parameters we obtain from our analyses.

In our method, the shape of the cornea in 3D microCT models is used to calculate the local inter-facet (IF) angle and facet diameter (indications of visual resolution and sensitivity, respectively), as well as the extent of its corneal projection (CP) as an approximation of the field of view (Taylor et al. 2019). These data can be displayed in world-based coordinates, which facilitates direct comparisons between eyes with different morphologies or sizes and in different preservation states. Because our method is non-destructive and is based only on the corneal structure, it is suitable to perform on fragile samples such as three-dimensionally preserved fossils or museum specimens, enabling the use of historical material for investigations into the factors that drive and shape the visual evolution of compound eyes.

We demonstrate the versatility of our analysis technique by examining the visual specializations of fungus gnats from different habitats and different evolutionary time periods. These insects are interesting models for studying the evolution of visual specialisations because, despite having eyes that comprise just a few hundred lenses and brains the size of a grain of salt, they are forest-dwellers, capable of using vision to guide behaviour in some of the most visually complex habitats on earth (Evenhuis 2006). Species of fungus gnats inhabit a broad range of forest types, from the arctic to the tropics, and their frequent occurrence in amber shows the tribe has thrived for at least 50 million years (Kraemer & Evenhuis 2008). We focus our study on females from three species of the cosmopolitan Orfeliini tribe, 1) an ancient species (Orfeliini sp.; Fig. S1) from the tropical rainforests of Scandinavia ~30 million years ago, 2) an extant species from the tropical rainforests of Africa (*Rutylapa* sp.), and 3) an extant species from the temperate forests of modern Scandinavia (*Neoplatyura modesta*). While both Scandinavian gnats are from similar geographic locations, they represent species from different habitat types, as forests in the Baltic region have thinned substantially since the Eocene (von Tschirnhaus & Hoffeins 2009). This study design exemplifies how our technique facilitates visual comparisons between extant species from different locations/habitats, and with extinct species from different points in environmental history from those same locations. Such comparisons are necessary if we are to understand the evolution of eye specializations in response to changing habitats (e.g. Scales & Butler 2016). For the continued evolutionary success of these diminutive Diptera, each species must be capable of acquiring the visual information necessary to control behaviour through its particular habitat. Yet, the eye specializations evolved for one habitat are unlikely to be optimal for other visual environments, due to differences in the density of trees, light intensity, and canopy cover. *Based on this consideration, we hypothesize that the visual properties of the fungus gnats from tropical habitats would be the most similar, despite the distance between their geographic and temporal distributions.* Specifically, given that tropical forests are relatively dark due to their dense canopies, we expected that tropical gnats could have specializations that improve their optical sensitivity and that these may not be possessed by gnats that inhabit brighter temperate forests. Although the focus on rare specimens naturally limits the scope of this work, we are nonetheless able to demonstrate the utility of our methodology for analysing the corneal structure of historical specimens by exploring whether the miniature eyes of gnats could have evolved *different* visual specializations depending on their specific visual habitat and if so, what are they?

## Methods and materials

### Animals

Each of the three fungus gnat species were preserved in a different way – dried, in alcohol, or as an amber endocast (Table S1). We only selected females for our analysis to minimize the effect of sexual dimorphism, as the males of many Diptera species have evolved visual specializations related to pursuit flights during courtship (Collett & Land 1975). The Entomology Collection at the Department of Biology, Lund University provided dried and pinned samples of *Neoplatyura modesta*, the Natural History Museum at the University of Oslo provided samples of *Rutylapa* sp. preserved in ethanol, while the Department of Geology at Lund University provided a piece of amber containing inclusions of Orfeliini sp. (Fig. S1). We found these samples varied in head width by approximately 20% (Table S1). Although shrinkage can occur in dried and ethanol preserved insect specimens (Friedrich et al. 2014), this is least apparent in the exoskeleton and a previous comparison between sample preservation methods indicated that ethanol fixation and drying do not obviously change the 3D appearance of Dipteran eyes (providing warping or collapse of the cornea is not present) (Sombke et al. 2015, personal observations).

### X-ray microtomography

X-ray microtomography (microCT) was performed on the head of each sample using a Zeiss XRM520 at the 4D Imaging Lab at Lund University. For each data set, X-ray projections were obtained over 360° (for specific scanning parameters see Table S2) and reconstructed into 3D volumes with 1*μ*m isotropic resolution. Due to their different preservation states, each sample was mounted in the tomograph using a different method, as outlined below.

- The pin of the dried sample was clamped in a pin-vice and mounted directly in the tomograph and was oriented such the pin did not occlude the projection of the gnat head on the detector.
- The alcohol-preserved sample was placed at the bottom of a 0.5 mL microcentrifuge tube (Eppendorf) and partially covered with a small amount of 70% ethanol (the surface tension of the liquid held the gnat in place). A ball of Parafilm was then used to fill the remaining volume of the tube and prevent evaporation of the ethanol. The initial tube was placed partially within a larger 2 mL microcentrifuge tube and secured with Parafilm, and the later was clamped in a pin-vice and mounted in the tomograph.
- The amber block was hot-glued to a ‘drawing pin’ that was clamped in a pin-vice and mounted in the tomograph. A lower resolution scan was initially conducted on the block to identify the location of several inclusions, with the head of one selected for the 1*μ*m resolution scans.

### General analysis procedure

The volume rendering from microCT scans allowed the facets of the compound eye to be identified (Fig. 1Ai) and a digital representation of the entire head surface to be computed (Fig. 1Aii) in Amira (FEI). The border of the left eye (Fig. 1Aiii) and each of its facets (Fig. 1Aiv) were then manually labelled by selecting paths across the surface. This isolated the surface of every individual facet on the left eye. The right eye was represented by mirroring the surface of the left eye (Fig. 1Av). For additional information on these steps, see the ‘Detailed surface analysis procedure’ section below.

**Figure 1:**
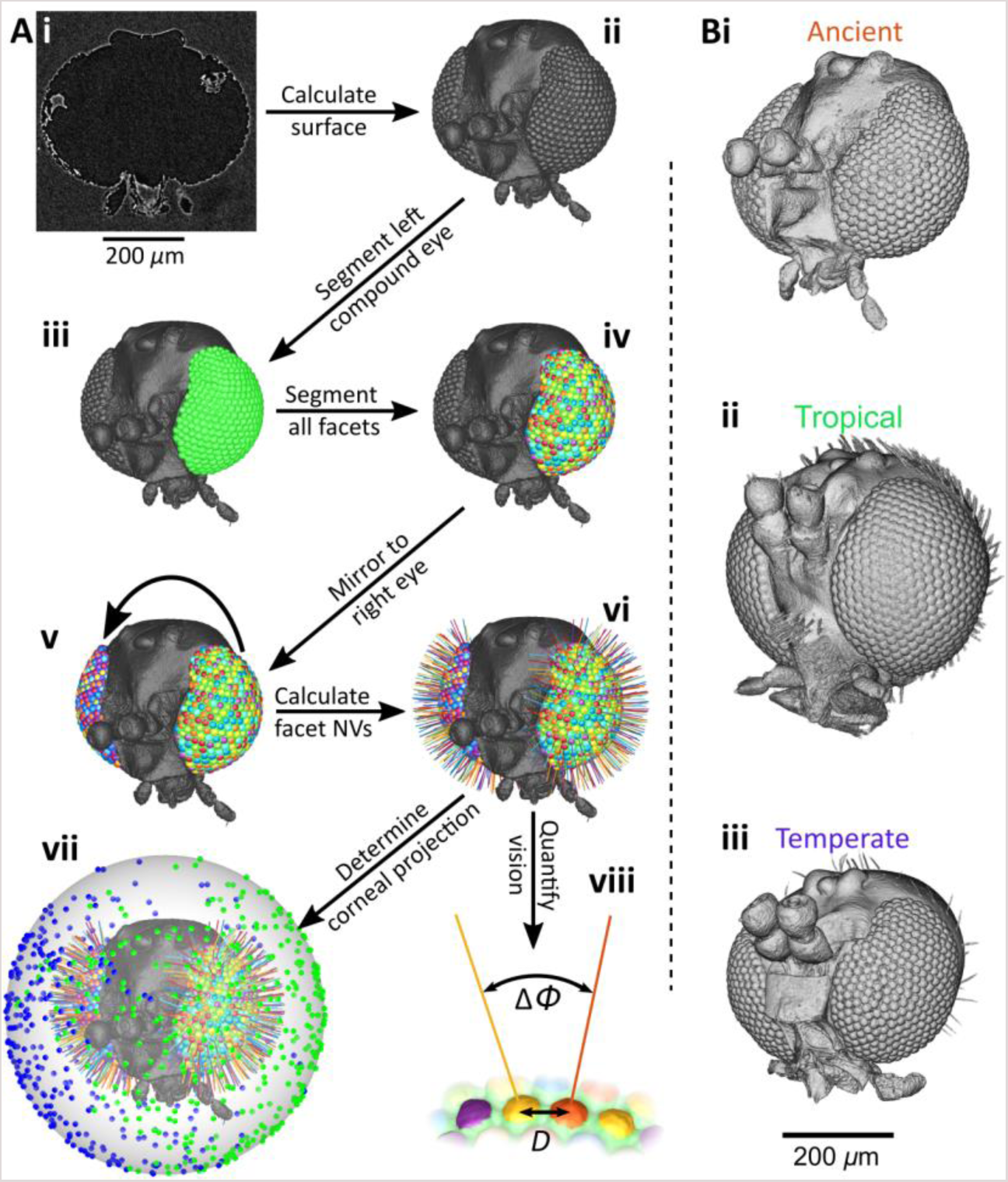
(**A**) Workflow to quantify the visual parameters of an insect cornea from X-ray microtomography. (**i**) Virtual section through a reconstructed volume showing the head of an Eocene fungus gnat from an amber endocast. (**ii**) The reconstructed exterior surface of the gnat’s head. (**iii**) The segmented left compound eye, and (**iv**) all of its segmented corneal facets. (**v**) The left eye mirrored to the right-hand side of the head, and (**vi**) the calculated normal vector (NV) from all facet surfaces. (**vii**) The NVs are projected onto a sphere to determine their viewing directions and visual fields (green – left eye, blue – right eye). (**viii**) Visual parameters can be calculated across the eye: facet diameter (*D*) and IF angle (Δ*Φ*). (**B**) Volume renderings of each gnat imaged for this study. (**i**) Endocast of Orfeliini sp. in Eocene Baltic amber; (**ii**) alcohol-preserved head of *Rutylapa* sp.; and (**iii**) dried head of *N. modesta*. The scale bar underneath Biii applies to all heads in B.

The normal vector (NV) of each facet (Fig.1Avi) and its viewing direction relative to the head (Fig.1Avii) were calculated from the surfaces after importing them into Matlab (Mathworks). The limits of the visual field of each eye were calculated from the NVs of their outermost facets. The complete visual field and binocular overlap was determined by combining the limits of both eyes (Fig. 1Avii). Facet diameters were calculated by finding the average distances between the centre of each facet and its neighbours (Fig. 1Aviii) and inter-facet (IF) angles were calculated by finding the average angle between the NV of a facet and its neighbours (Fig. 1Aviii) (Land 1997). Diagrams of how facet diameter and IF angle vary over the visual field were generated from the visual axes and properties calculated for each gnat. A voronoi diagram was drawn around the visual axes of each eye, and the cells were coloured according to the local facet diameter (Fig. 2A-C:i) or IF angle (Fig. 2A-C:ii), which could also be depicted by colouring the individual facets of the compound eye (Fig. 2A-C insets). This is an adaption of the analysis procedure we recently developed (Taylor et al. 2019), with a key conceptual difference being that here calculations are performed on individually labelled facets, rather than being interpolated between sparse labels. A simulation of how each gnat may have viewed a forest scene was generated from a panoramic forest image, considered to represent the type of scene that these insects may have viewed. In the simulation, we assumed that each facet accepted light over an angle equal to its IF angle, and then coloured the voronoi cells based on the weighted average of the intensity value of the pixels that lay within this angular range (Fig. 3). For additional information on these steps, see the ‘Detailed computational analysis procedure’ section in the supplemental material.

**Figure 2:**
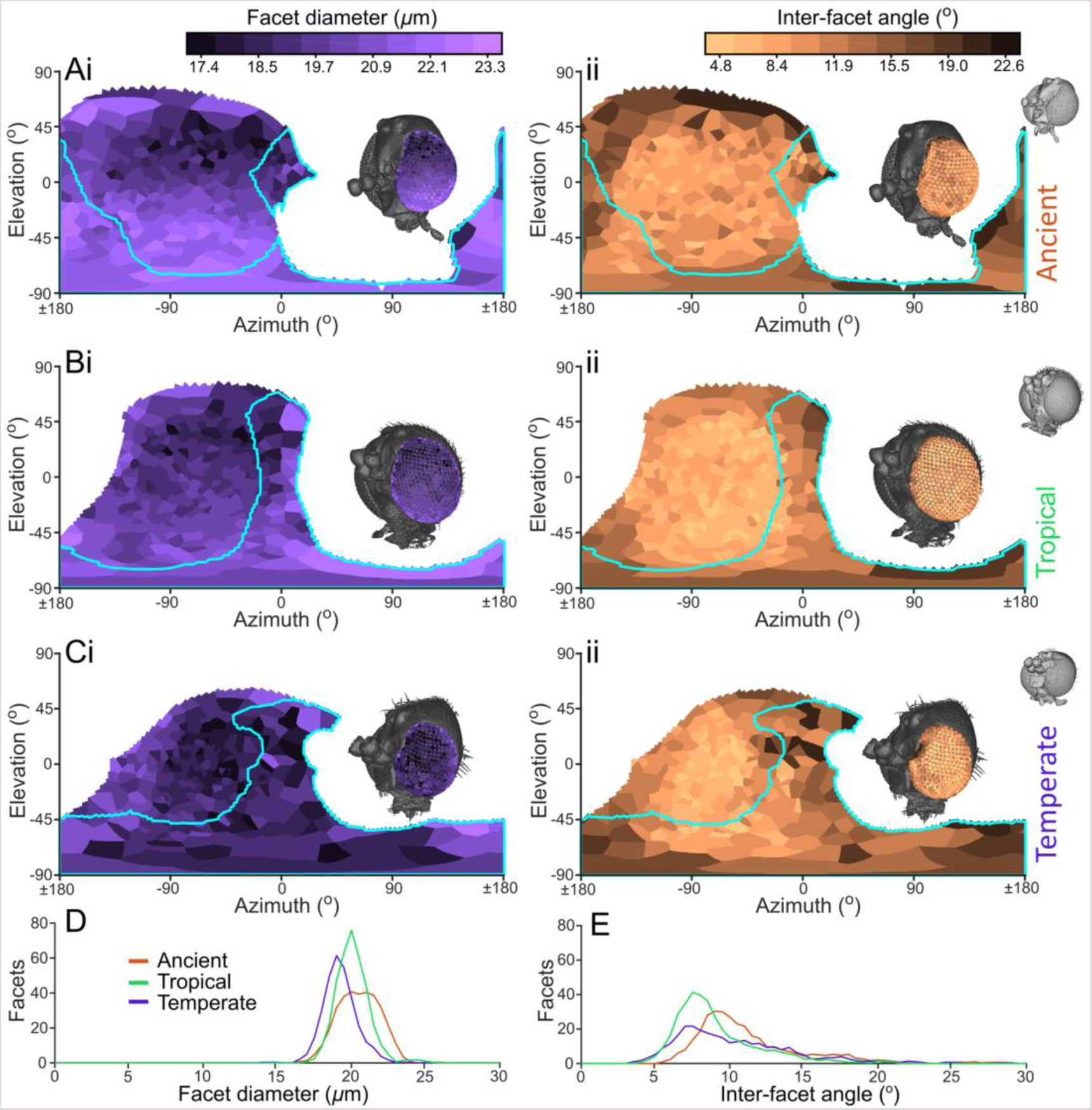
Visual parameters of fungus gnat eyes for (**A**) the ancient gnat, (**B**) the extant tropical gnat, and (**C**) the extant temperate gnat, with maps of (**i**) the facet diameter and (**ii**) the inter-facet (IF) angle, as they are projected onto the visual world. These parameters are also shown directly on the corneal facets in the inset of each panel. Note that a lower angle indicates higher resolution vision. Each gnat is facing towards 0° azimuth, and the cyan lines indicate the limit of binocular corneal projection (CP). The frequency distribution of facet diameter and IF angle, for each individual are shown in (**D**) and (**E**), respectively.

**Figure 3:**
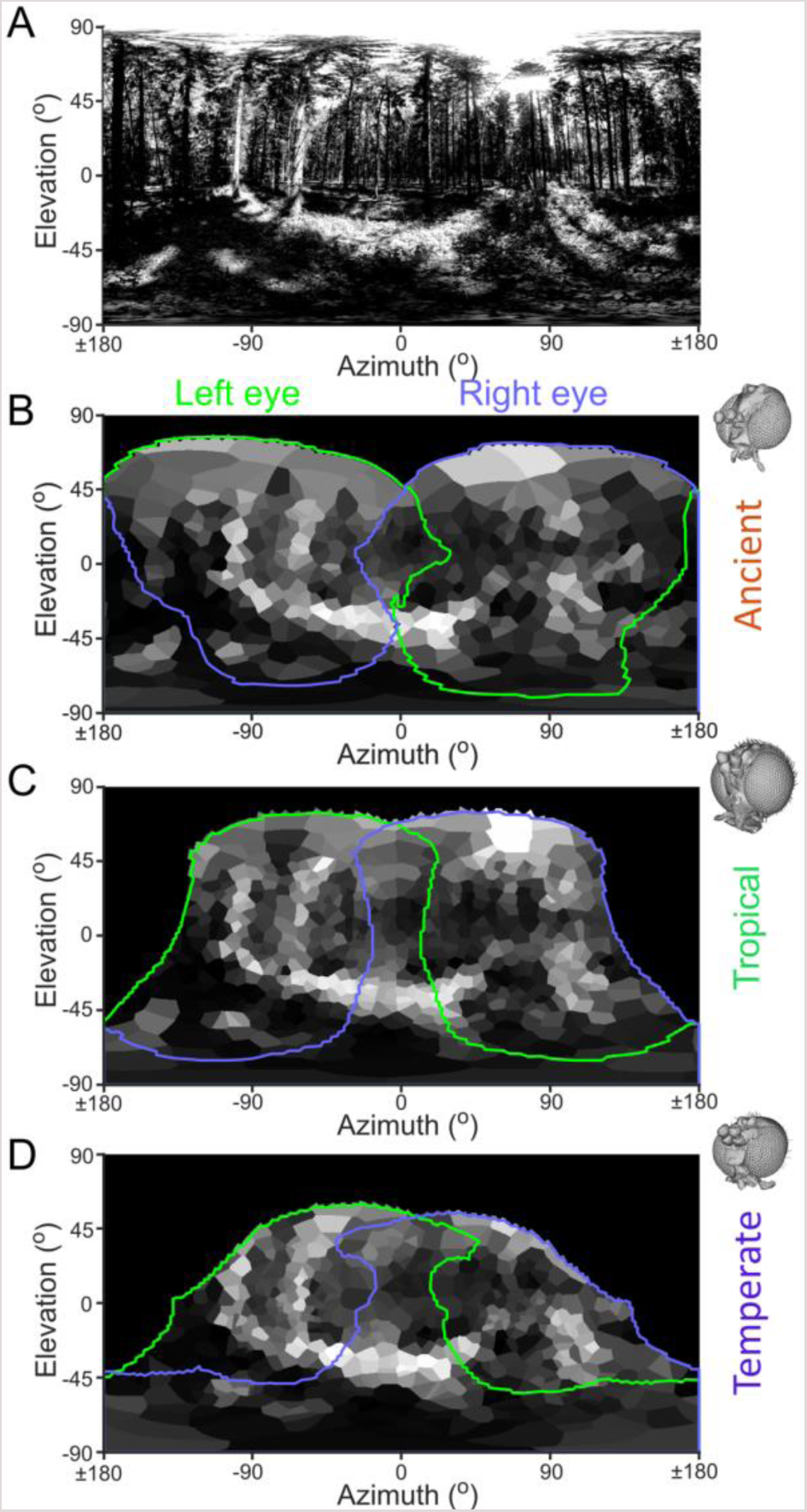
A forest viewed through the eyes of different gnat species. (**A**) An equirectangular projection of a 360° panoramic image of a European forest in summer (image: istock.com/Bestgreenscreen). (**B**) Simulated view of the scene with data quantified from the eyes of the ancient gnat, and likewise, from the eyes of the extant tropical (**C**) and the temperate (**D**) gnats. The green and purple lines denote the limits of the corneal projection (CP, approximating the full field of view) for the left and right eyes, respectively. The simulation accounted for the variation in inter-facet (IF) angle (but *not* any spatial variation in optical sensitivity) across each eye’s CP.

## Results

Our analysis technique uses laboratory microCT to image the 3D structure of the heads and eyes of insects (Fig. 1Ai, ii). We then calculate the visual properties of the eyes from the structure, shape, and size of the individual corneal facets (Fig. 1Aiii-v). The apposition compound eyes of arthropods are composed of many facets, also known as ommatidial units. Each ommatidium comprises a corneal lens, a light-guiding cone and an underlying group of photoreceptors (Land & Nilsson 2012). In the specimens we analysed, the cornea was the only intact part of the eye, but this could nonetheless be used to determine facet diameter, the local inter-facet (IF) angle, and the full corneal projection (CP). The diameter of the facets provides an indication of the amount of light that is focused onto the underlying photoreceptors and can thus be used to approximate optical sensitivity – larger facets capture additional light in proportion to the square of their diameter and increase optical sensitivity considerably. The IF angle (calculated using the surface normal vector, NV, of each facet, Fig. 1viii), can be used to approximate the inter-ommatidial angle, which in turn can be used to predict visual resolution, because each lens focuses light from a specific region of space (its viewing direction) centred about its optical axis. The smaller the angle between the optical axes of adjacent facets, the greater the visual resolution is likely to be (Land 1997). In turn, the NVs can be used to calculate the angular projection of the cornea into the world (the CP), which approximates the entire eye’s field of view (Fig. 1Avi, vii). It is important to note that our measures provide only approximations because we are limited to taking measurements from the cornea. For example, sensitivity is also affected by the receptor dimensions and its optical acceptance angle (variables that cannot usually be measured in natural preserved specimens due to degradation of internal structures) (Land 1997) and discrepancies between the IF and the inter-ommatidial angle can differ if the cone and rhabdom that underlie the lens are skewed relative to its NV, (although this is usually only seen at the edges of the eye) (Stavenga 1979).

To explore whether the visual systems of fungus gnats have changed both across habitat type and across time, we applied our technique to three species (Orfeliini sp. – ancient, tropical, Fig. 1Bi; *Rutylapa* sp. – extant, tropical, Fig. 1Bii; *N. modesta* – extant, temperate, Fig. 1Biii). The exterior corneal surfaces reconstructed from microCT showed that, superficially, the ancient gnat appears most similar to the extant temperate species (Fig. 1B). While its head and eye size lay between those of the two extant gnats (Table S1), the ancient gnat had both larger facet diameters (Fig. 2D) and IF angles (Fig. 2E), and also a larger CP – indicating a larger field of view – than either extant species (Table S1). Interestingly, the CPs of all species had a binocular overlap that is directed ventrally and frontally (Fig. 2A-C). Despite the gnats’ small eyes, it is evident that they also possess regional visual specializations. For example, the IF angles vary across the visual field of all gnats. The lowest IF angles (indicating higher resolution) are directed laterally and view their monocular CP (Fig 2A-C:ii). Furthermore, a ventral-to-dorsal reduction in facet diameter is visible across the CPs of both tropical specimens (Fig. 2A,B:i), but this is not apparent in the temperate gnat (Fig. 2C:i). A simulation of the spatial sampling of both eyes of each gnat indicates that all three should obtain coarse, but nonetheless distinct, spatial information from the features in a forest scene (Fig. 3).

## Discussion

Vision is essential for guiding many animal behaviours, but we know little about the intrinsic and extrinsic factors driving the evolution of visual systems or their adaptation to specific visual environments. This knowledge gap is partly due to the limitations of current methodologies for quantifying eye specializations, which are best applied to fresh or preserved tissue (Stavenga 1979, Ullmann et al. 2012), thereby preventing investigations of rare, naturally preserved, or ancient specimens. To overcome this, we have developed a new method for uncovering the visual world of arthropods using non-destructive microCT methods (Fig. 1Ai) that can be applied equally well to the cornea of fresh, dried, or even fossilized compound eyes. We demonstrated this method by quantifying the vision of fungus gnats’ tiny eyes.

While not readily apparent from their external morphology (Fig. 1B), we can identify visual specializations from our analyses of fungus gnat species from different locations, habitats and historical time periods. In particular, we identified two visual specializations *common* to all three of the Orfeliini species investigated. The first is that their highest resolution vision is directed laterally, and lies outside of their binocular field (Fig. 2A-C:i). This is in contrast to larger Diptera (including *Drosophila*), where the highest resolution is directed frontally (Heisenberg & Wolf 1984, Land & Eckert 1985, Straw et al. 2006). The second specialization common to the species investigated here is that they have a large binocular overlap that is primarily directed ventrally and somewhat frontally (Fig. 3). Interestingly, the extent of binocular overlap in larger (female) flying insects is typically smaller and approximately ventrally-to-dorsally symmetric (Beersma et al. 1977, Seidl & Kaiser 1981, Merry et al. 2006). We propose that ventral binocularity is a previously unconsidered adaption for small, forest-dwelling insects that could allow them to improve sensitivity by integrating the noisy signal of shadowed objects on the ground between both their eyes. Our method also allowed us to identify that gnats have distinct *differences* between their vision, which may represent habitat-specific specializations. We find that both the extant and ancient tropical gnat species have larger, more dorsally-directed total corneal projections (CPs) than the temperate gnat (Fig. 3). Tropical species also had slightly larger facet diameters than temperate species (Fig. 2D & Table S1), indicating a global increase in sensitivity. Additionally, while the facet diameters remain relatively constant across the temperate gnat’s eyes, those in the tropical gnats have a clear negative relationship to elevation, as their ventral facets are larger than those facing dorsally (Fig. 2A,B:i). These larger ventral facets appear to represent a regional visual specialization that would likely improve the relative sensitivity of the ommatidia viewing the forest floor. An analogous increase in eye ‘regionalization’ is also found among extant damselfly species that live in dark or visually complex habitats (Scales & Butler 2016). The increased sensitivity provided by the combination of both larger ventral facets and binocular overlap could substantially enhance the reliability with which tropical gnats can discriminate objects beneath them and therefore reduce the need to develop larger eyes. Taken together, our findings support our hypothesis that the tropical gnat species have habitat-specific specializations, primarily in the form of locally-increased optical sensitivity, compared to their temperate relatives.

Our results demonstrate that this advance on our analysis method can be used to identify visual specializations present in species from different times and environments, and will enable investigations into the visual systems of rare or extinct species in various states of preservation. While the limited (but unavoidable) sampling of this case-study provides only tentative support for our habitat-hypothesis, these preliminary results provide inspiration and justification for a larger, more detailed study considering variation both within-species and within-habitat groups while controlling for phylogenetic effects. A distinct advantage of our method is that it facilitates direct comparisons of vision between species. Unlike existing approaches, the method enables the visual properties of each insect to be presented in a common, world-based coordinate frame, even if there are substantial differences between the visual morphology or size of the eyes (Taylor et al. 2019). Additionally, the ability of this technique to simulate vision through an arthropod’s eyes provides insights into how it views a particular scene which, given the substantial differences between arthropod vision and our own, is a valuable tool for interpreting visual parameters and developing hypotheses about the function of specific visual adaptations (Fig. 3). For instance, despite being weak fliers, fungus gnats are effective pollinators (Mesler et al. 1980). Is it possible that the eyes of tropical species allow them to find flowers more easily in a rainforest than the eyes of a temperate species would? Gaining knowledge about the visual capabilities of fungus gnats and being able to simulate their visual world is a valuable starting point for designing behavioural assays to conclusively answer such a question – which may become more important given the current drastic declines of flying insect biomass due to habitat change (Hallmann et al. 2017).

The corneas of insect eyes can remain well-preserved in amber dating back to the Early Cretaceous (Poinar & Milki 2001) and we expect that there are many further opportunities to use our methodology to study the circumstances through which specific visual traits evolved in other taxa. For example, prominent acute zones with regions of flattened and enlarged facets have evolved to accompany pursuit behaviour in many insect species. Identification of such regional specializations in the fossil record could, for instance, be used to investigate the evolutionary origins of pursuit mating strategies in male bees (Streinzer & Spaethe 2014). Fine retinal structures can be identified in our alcohol-preserved sample (Fig. S2) and are also likely to be visible in images of exceptionally-preserved amber inclusions (Tanaka et al. 2009). MicroCT has been used to describe the morphology of insect inclusions for taxonomic purposes (Lak et al. 2008, Greco et al. 2011) and data obtained for systematics could also be reused to describe a specimen’s vision. Applying our methodology to the substantial range of dried specimens collected over the last two centuries would also facilitate investigations into the influence of recent anthropogenic disruptions on the evolution of invertebrate visual systems. For example, it could be used to examine whether the increase in light pollution over the last century has placed any selection pressure for specific visual traits on the eyes of nocturnal urban insects (Hölker et al. 2010, Knop et al. 2017). Testing these, and many other hypotheses related to the factors driving visual evolution will be possible by applying this technique to analyse the well-preserved compound eyes of the arthropod specimens available in museum collections.

## Acknowledgments

We would like to thank Rune Bygebjerg from the Department of Biology at Lund University and Gier Søli from the Natural History Museum at the University of Oslo for supplying the extant samples used in this study, and Vladimir Blagoderov from the London Natural History Museum for assisting with the identification of the amber embedded specimen. Additionally, we are most indebted to Viktor Håkansson for his meticulous segmentation of the data. Gavin Taylor is thankful to have received a stipend from Carl Tryggers Stiftelse (CTS15:38) and an endowment from the Royal Physiographic Society of Lund, while Emily Baird gratefully acknowledges financial support from the Air Force Office of Scientific Research (FA8655-12-1-2136), the Swedish Research Council (2014-4762), and the Lund University Natural Sciences Faculty.

## Author Contributions

Conceptualization, G.J.T., J.A.G., and E.B.; Methodology, G.J.T, J.A.G., S.A.H., and E.B.; Software, G.J.T.; Investigation, G.J.T. and E.B.; Formal Analysis, G.J.T. and S.A.H.; Writing – Original Draft, G.J.T. and E.B.; Writing – Review & Editing, G.J.T., J.A.G., S.A.H., and E.B.; Visualization, G.J.T.; Funding Acquisition, G.J.T. and E.B.; Resources, J.A.G.; Supervision, G.J.T. and E.B.

## Supplemental material

### Detailed surface analysis procedure

Each image stack from microCT was imported into Amira (v6.2, FEI), within which an interactive workflow was developed that allowed us to create a surface defining the head of the scanned insect, and then to delineate the border of the left eye plus each of its facets. After labelling the facets, these data were mirrored to the right eye, and the surface data and mirror transform were then exported to Matlab for further computational analysis (see the following section). In the following descriptions, windows within the Amira console are indicated in *italics*, Editor windows (accessible from the *Properties View* of a module) are indicated by <u>underlining</u>, and further modules, tools, and options are indicated in ‘parenthesis’.

The head surface was created by first labelling the voxels within the head volume, and then fitting a surface around their border, as described in the following. The ‘Threshold’ tool in the *Segmentation View* was used to select and label the voxels that were part of a gnat (which were lower intensity than the surrounding material for the amber-embedded specimen, but higher in the other two cases). The ‘3D Lasso’ tool was then used to select and remove excess labelled material, such that only the fungus gnat’s head capsule remained labelled. Following this, a surface was created around the labels using the ‘Generate Surface’ (a ‘smoothing extent’ of 2 was used when generating the surface) and ‘View Surface’ modules in the *Project View*. Sometimes the surface created for the eyes appeared to be poorly formed, particularly for the amber-embedded and alcohol-preserved samples. In the former case, this was because decayed material from the gnat coated the inside of the endocast, and often had higher absorption than the amber. In the alcohol-preserved case, it was because of relatively low absorption contrast between the cuticle and the alcohol. In both cases, the ‘Brush’ tool was used to manually correct the label field of 2D slices in the *Segmentation Editor*, to ensure that individual facet surfaces of the left eye appeared smooth in the generated surface and, additionally, that the surface of the eye was closed, with no gaps present between facets. The scan of the dried gnat had high contrast between the cuticle and air and the initial surface that was generated did not require such corrections.

After completing any required corrections, a final surface was created and a ‘Create Surface Geodesic Path’ module was attached to it (all remaining steps are performed in the *Project View*). In the <u>Surface Path</u> <u>Editor</u>, the connector was set to ‘dijkstra’, and the control points to ‘vertex’. The <u>Surface Path Editor</u> allows points to be selected around the border of the left compound eye, which were linked by paths taking the shortest route across the surface between sequential points. To isolate this area from the remainder of the surface, after closing the path around the eye, the ‘Patchify Surface’ was selected. The <u>Surface Editor</u> was then used to selectively display only the compound eye patch, after which the ‘Extract Surface’ module was used to create a new surface only including the compound eye. A second ‘Create Surface Geodesic Path’ module was then attached to the isolated corneal surface and paths were traced in a circle around the base of every facet on the compound eye. Approximately 60 control points were placed per facet and closing a path upon completion allowed a new path to be started for the next facet. Note that each fungus gnat facet has a dome shaped cornea; many other insects have flatter, hexagonal lenses and placing a control point at each of the six corner points of a facet would probably be sufficient in such cases. The right eyes of gnats were not individually segmented, but we counted the number of facets by using another geodesic path to place control points at the centre of each facet. This indicated that the gnat eyes were approximately symmetrical, with at most a 2.4% difference in facet number between the eyes (Table S1).

Each head surface was then aligned using the <u>Transform Editor</u>, by positioning and aligning it such that it faced forwards in the default XY image in the *Project view*. The head was aligned in its frontal (yaw) and axial (roll) planes based on its symmetry. The pitch (sagittal plane) of the head could not be aligned by symmetry, but was positioned such that each head appeared to have a similar, forwards facing, orientation (Fig. 2A-C) (Taylor et al. 2019). While the orientation of the head may vary during flight this provides a consistent reference frame between insects. After aligning the head, the resulting rigid transformation matrix was copied to both the isolated eye surface and the paths around each facet. While the right eye was not separately labelled, we determined the transformation required to mirror the labelled left eye such that it was aligned with the position of the right eye. To mirror the left eye, we used the ‘Scan surface to volume’ module on the left eye surface to create a volume in which the voxels corresponding to the surface were labelled. The ‘Resample transformed image’ module was then applied to this volume, which created a second volume, the axes of which were orthogonal to the world coordinate frame. This volume could then be mirrored around the sagittal plane of the head by using the ‘Flip X’ button in the <u>Crop Editor</u>, and a ‘Volume rendering’ module was used to visualize the position of the mirrored eye. Finally, the <u>Transform Editor</u> was used to manually adjust the alignment of the mirrored right eye such that it visually matched the position of the actual right eye on the surface of the head as closely as possible.

After completing the above-described processing workflow in Amira, it was necessary to export data for use in Matlab. Both the head surface and eye surface were exported as ‘stl’ files in little endian format. The paths surrounding each facet were exported as ‘Amira lineset’ files in asci format. To facilitate importing paths into Matlab, it was necessary to split the lineset file into two parts using a text editing program. Each file contains data under two headings; @1 is a list of indices indicating the vertices to use for each control point in an individual path (−1 indicates that start of a new path), @2 is a list of 3D vertices coordinates referenced by the indices. It is necessary to copy the indices and vertices into individual text files. Finally, the 4×4 transform matrices for the head and the right eye were recorded by using the ‘GetTransform’ command in the *TCL Console*, and the ‘Max index’ and the ‘Min coord’ values of the right eye volume were recorded from <u>Crop Editor</u>.

**Upon acceptance of this manuscript: We will submit the original volumes and head surfaces from each sample to the MorphoSource depository and include their DOIs in the manuscript. We will also submit an example Amira project showing the result of the complete workflow described above to Data Dryad. Finally**, **we will submit the files exported from Amira to Data Dryad so that the full results of the Matlab scripts described below can be replicated.**

### Detail computational analysis procedure

The data exported by Amira was imported into Matlab and analysed using a series of custom written scripts to quantify the visual properties of gnat eyes and perform a visual simulation. The analysis procedure involves applying the rigid transformations to the imported surfaces and identifying the facet surfaces from the path points. The corneal normal vector (NV) from each facet was calculated, which was used to quantify the eye’s visual parameters and fields of view (Taylor et al. 2019). Various plotting options were implemented for the data, including performing a visual simulation of the insect’s perspective of a scene. This analysis was mostly automated, and here we describe the analysis steps performed by the script. Variable names that the user may wish to modify are indicated in ‘parenthesis’, and influence data importing and various display options as outlined in the initial ‘Configuration’ section of the supplemental ‘AnalyseCorneaMain.m’ file.

Data were initially imported into Matlab based on the user specified files in the initial portion of the ‘Configuration’ section. The exported transformations were applied to the surfaces such that they were represented in the same orientation as seen in Amira. The paths were initially formatted as a single list of points, but we separated this into an individual list of vertices for each facet’s border. After this, the coordinates on the eye surface enclosed by each border path were selected as the corneal surface for that facet. The centre point of each facet was selected as the surface point that lay closest to average coordinate of the border path (in a 2D projection), and then the NV of the cornea was calculated from this point. This calculation involved fitting a second order polynomial surface to the corneal surface vertices, and calculating the normal direction from the derivative at the central point (Taylor et al. 2016, Taylor et al. 2019). The NV was taken to indicate the optical axis of the facet. The same procedure was performed for the right eye using the facets mirrored from the left eye.

The neighbours of each facet were determined based on the shortest distances between its centre and those of each other facet (each facet linked to six neighbours, unless it was on the border of the eye). Having established the neighbours of each facet, the inter-facet (IF) angle of a given facet was calculated as the average angle between the normal from its centre and those of its neighbours and the facet diameter was calculated as the average distance from its centre point to those of its neighbours. The eye parameter, was calculated by multiplying each facet’s IF angle by its diameter (Snyder 1979). Note that the calculating the eye parameter assumed that the inter-ommatidial angle equals the computed IF angle, which may not be true and limits the accuracy with which we can report this parameter. It is possible to perform a local smoothing operation on the data, whereby the NVs and diameters of each facet are averaged with those of their neighbours. Smoothing may be useful if the visual parameters of an eye vary irregularly and can be adjusted using the ‘smoothingFactor’ variable (we did not smooth the results shown in this study).

To approximate the field of view of each eye, we determined its corneal projection (CP). We did this by first calculating the intersection of each NV on a distant sphere. We then discretized the sphere into many equally-spaced points and determined which of those points lay within a boundary enclosing the NV intersections. The points within the CP were determined in this manner for each eye individually. The union of these two sets of points provided the total CP, while the intersection indicated the portion of the CP with binocular overlap. The borders around each region were also calculated for use in the later plotting steps; note that if multiple individual regions are present within a given CP (such as in the binocular CP, Fig. 2A), the border calculation may take a long time to compute. The visual sphere was also divided into individual regions for each facet. This was calculated either for the facets of the left eye only (Fig. 2) or for both eyes (Fig. 3) by calculating the borders of the voronoi cells (on the surface of the sphere) around the sphere intersection points of the facets. Although the voronoi cells covered the entire sphere, those on the borders were limited to the extent of the previously calculated CP.

Different methods were implemented to plot the data resulting from the previous analysis. The individual NVs and CPs were drawn onto a sphere surrounding a visualization of the head surface (Fig. S3A). The values of each calculated parameter were encoded using colour maps, and displayed either as a world-based representation by colouring the voronoi cells of the left eye (Fig. 2A-C), or as a head-based representation by colouring the corneal facets on the eye surface (Fig. 2A-C insets). While the voronoi cells were calculated on a sphere, we displayed them as 2D maps using an equirectangular projection, onto which either the binocular CP or the right eye CP could be displayed using the ‘plotBinoLine’ and the ‘plotRightEyeLine’ variables, respectively. It was also possible to draw contour lines of one parameter upon the map of another using the ‘dispContourOnInterOImage’ variable (for example contours of facet diameter on a map of IF angle, Fig. S3B). Facets viewing the binocular CP could also be indicated with markers on the head centric representation (using the ‘binoMarkerSize’ variable). All plots could be saved automatically by setting the ‘saveImages’ flag.

To display colour maps, the observed range of each calculated parameter was discretized into a number of equally spaced bins (set with ‘numColBins’ variable). By default, this discretization is performed for each insect, and different insect eyes had different ranges and receive different colour mappings. To compare between multiple insects, ‘AnalyseCorneaMain.m’ should first be run for each insect with the ‘saveData’ flag set. This will save the calculated parameters for each insect analysed and the ‘PlotGroupHistograms.m’ script (which requires similar variables to be set as in the main analysis file) was used to produce overlaid histograms for the parameters, and to compute the mean, standard deviation, and range of each parameter for each insect. The latter file also determined the total ranges of each calculated parameter between all data sets and used them to compute the appropriate discretization for all insects. In the original file, the ‘useGroupColBins’ flag should then be set, and re-running the main analysis file for each insect creates plots with a common colour mappings.

A simulation of an insect’s vision of a user-supplied panoramic image (Fig. 3) or a chequered sphere (Fig. S3D) could also be computed (choose by setting the ‘simUserIm’ flag). When using the chequered sphere, the number of chequers was set using the ‘numChequers’ variable (note that a higher number of chequers results in a lower wavelength and higher spatial frequency) and that chequers are mapped onto equirectangular image and do not represent equal angular areas on the world sphere (Fig. S3C). The simulation was performed using a direct RGB-to-grayscale conversion from the image provided as no knowledge of the spectral sensitivity of the insects was available. The world coordinates were determined for each pixel of the image being used, and we simulated the insect’s position at the centre of the world sphere. For each facet, we averaged the intensity of all points that lay less than one IF angle from its corneal axis and weighted this average by a Gaussian kernel with full width at half maximum (FWHM) equal to the IF angle of each facet. This assumes that the acceptance angle equals the IF angle for each facet (this proportion can be adjusted using the ‘deltaPScale’ variable). We then shaded the facet’s voronoi cell with the averaged grayscale value, and the simulation could either be performed for the left eye only or for both eyes by setting the ‘simBothEyes’ variable. The shaded cells were plotted on an equirectangular image in a similar manner to the colour maps described above, and the border of the fields of view could also be displayed.

**Upon acceptance of this manuscript: We will submit the Matlab files required to complete the analysis procedure described to GitHub. We will also submit the calculated visual properties to Data Dryad.**

## Supplemental tables

**Table S1:** Summary of additional parameters related to each fungus gnat specimen. Corneal projection (CP) is indicated as a percentage of the visual sphere viewed. Visual field extents are also provided as the minima and maxima angular limits of the total azimuthal range (at 0° elevation) and the total elevation range (at 0° azimuth), followed by the total angular extent in parenthesis (the CP behind the head is indicated with values from −90° to −270° in the elevation range). The mean ± one standard deviation is provided for the facet diameter, IF angle, and eye parameter, followed by the measured minima and maxima values in parenthesis.

| Species | *Neoplatyura modesta* | Orfeliini sp. | *Rutylapa* sp. |
| --- | --- | --- | --- |
| Habitat | Temperate forest | Tropical forest (Eocene) | Tropical forest |
| Collection location and year | Stenshuvud, Sweden (1977) | Lilla Beddinge, Sweden (1990) | West Usambaras Mountains, Tanzania (1990) |
| Preservation method | Dried | Endocast in amber | Alcohol |
| Left eye facet # | 303 | 333 | 367 |
| Right eye facet # | 296 | 335 | 373 |
| Head width ($\mu\text{m}$) | 452 | 519 | 547 |
| Left eye CP (%) | 48 | 55 | 46 |
| Total CP (%) | 69 | 96 | 78 |
| Binocular CP (%) | 27 | 17 | 15 |
| Total azimuth range (°) | -138 – 138 (276) | -180 – 180 (360) | -128 – 135 (263) |
| Total elevation range (°) | -131 – 55 (186) | -228 – 44 (268) | -140 – 66 (208) |
| Facet diameter, $D$ ($\mu\text{m}$) | $19.2 \pm 1.0$ (14.7 – 23.4) | $20.5 \pm 1.3$ (16.8 – 25.6) | $20.1 \pm 1.0$ (17.7 – 25.1) |
| Inter-facet (IF) angle, $\Delta\Phi$ (°) | $10.5 \pm 4.5$ (3.9 – 40.1) | $11.3 \pm 3.7$ (6.0 – 28.5) | $9.1 \pm 2.8$ (4.3 – 20.6) |
| Eye parameter, $P$ ($\mu\text{m}.\text{rad}$) | $3.6 \pm 1.6$ (1.3 – 14.1) | $4.1 \pm 1.4$ (1.9 – 12.5) | $3.2 \pm 1.1$ (1.4 – 9.0) |

**Table S2:** The specific scan settings for the Zeiss XRM520 tomograph that were used for imaging. The 4x objective was used for all samples.

| Species | *Neoplatyura modesta* | Orfeliini sp. | *Rutylapa* sp. |
| --- | --- | --- | --- |
| Source voltage (kV) | 80 | 80 | 50 |
| Source power (W) | 7 | 7 | 4 |
| Exposure time (s) | 2 | 10 | 10 |
| Projections (#) | 1,601 | 2,001 | 1,601 |
| Source to rotation axis distance (mm) | 15.0 | 21.0 | 14.0 |
| Rotation axis to detector distance (mm) | 86.3 | 120.9 | 80.0 |

## Supplemental Figures

**Figure S1:**
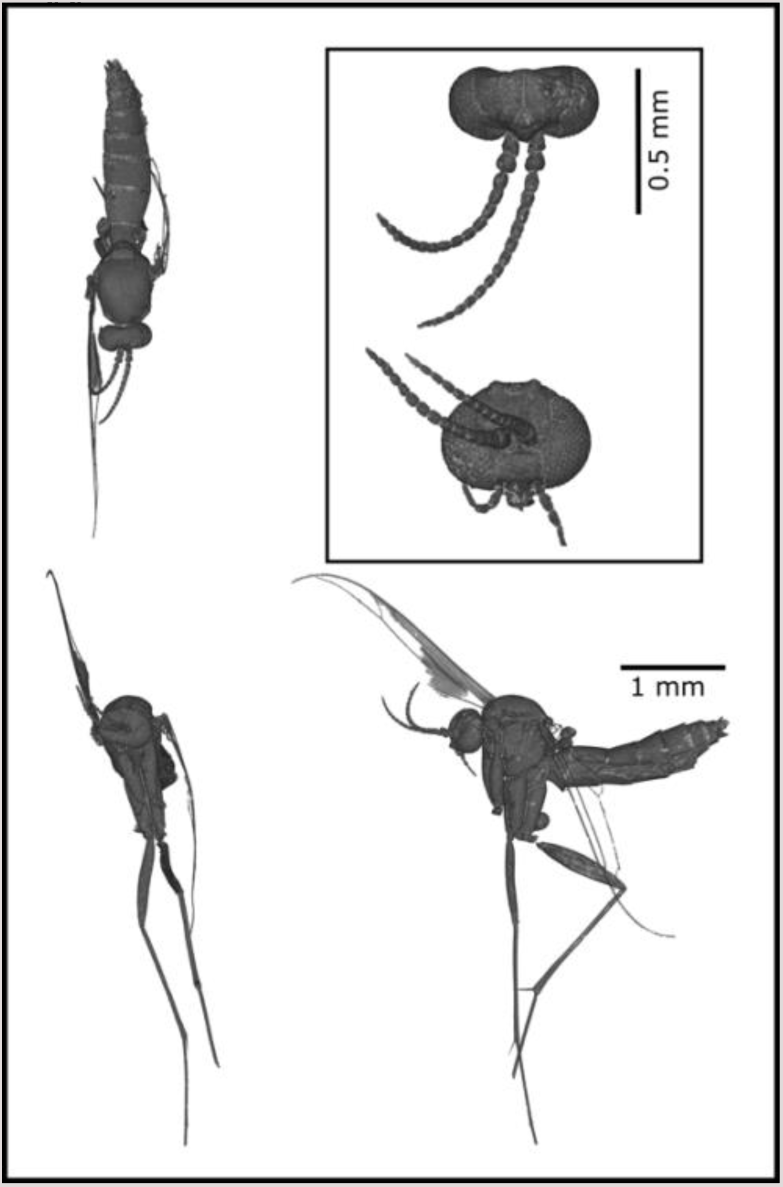
Habitus image created from the endocast of the female Eocene gnat specimen. Unfortunately, the degradation of the wings limited a full species identification of the sample, but the general morphological features place it in the tribe Orfeliini of the family Keroplatidae in the order Diptera.

**Figure S2:**
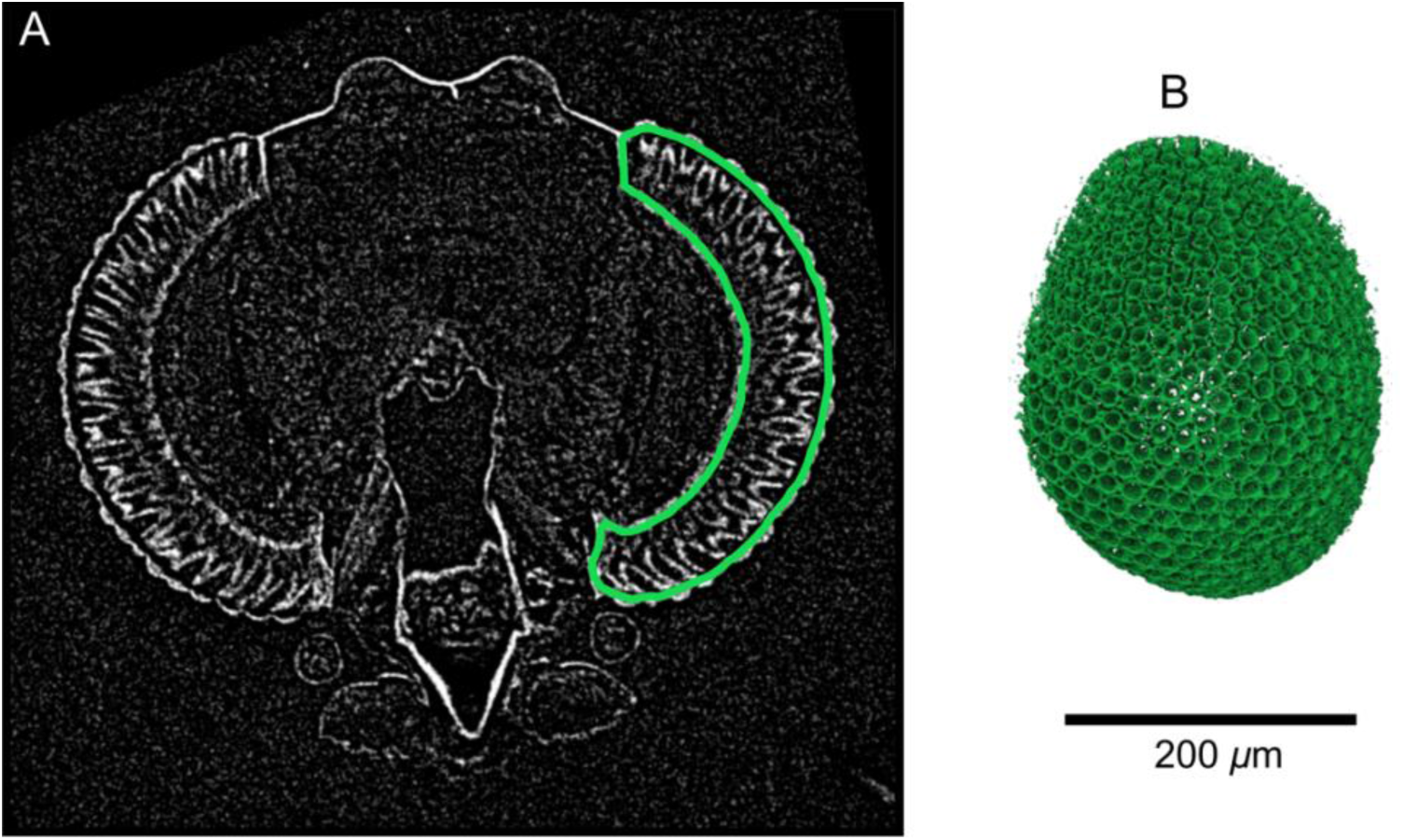
Internal structure of the alcohol-preserved *Rutylapa* sp. specimen. (**A**) A slice through the image volume shows that, in addition to the exterior cuticle, details within the retina of the eyes have high x-ray absorption. (B) We segmented the retinal volume of the left eye (enclosed by the green line in A), and used volume rendering to visualize the 3D retina. A conical x-ray absorbing structure is visible for each ommatidia, which may represent the secondary pigment cells surrounding the photoreceptors. The scale bar applies to both A and B.

**Figure S3:**
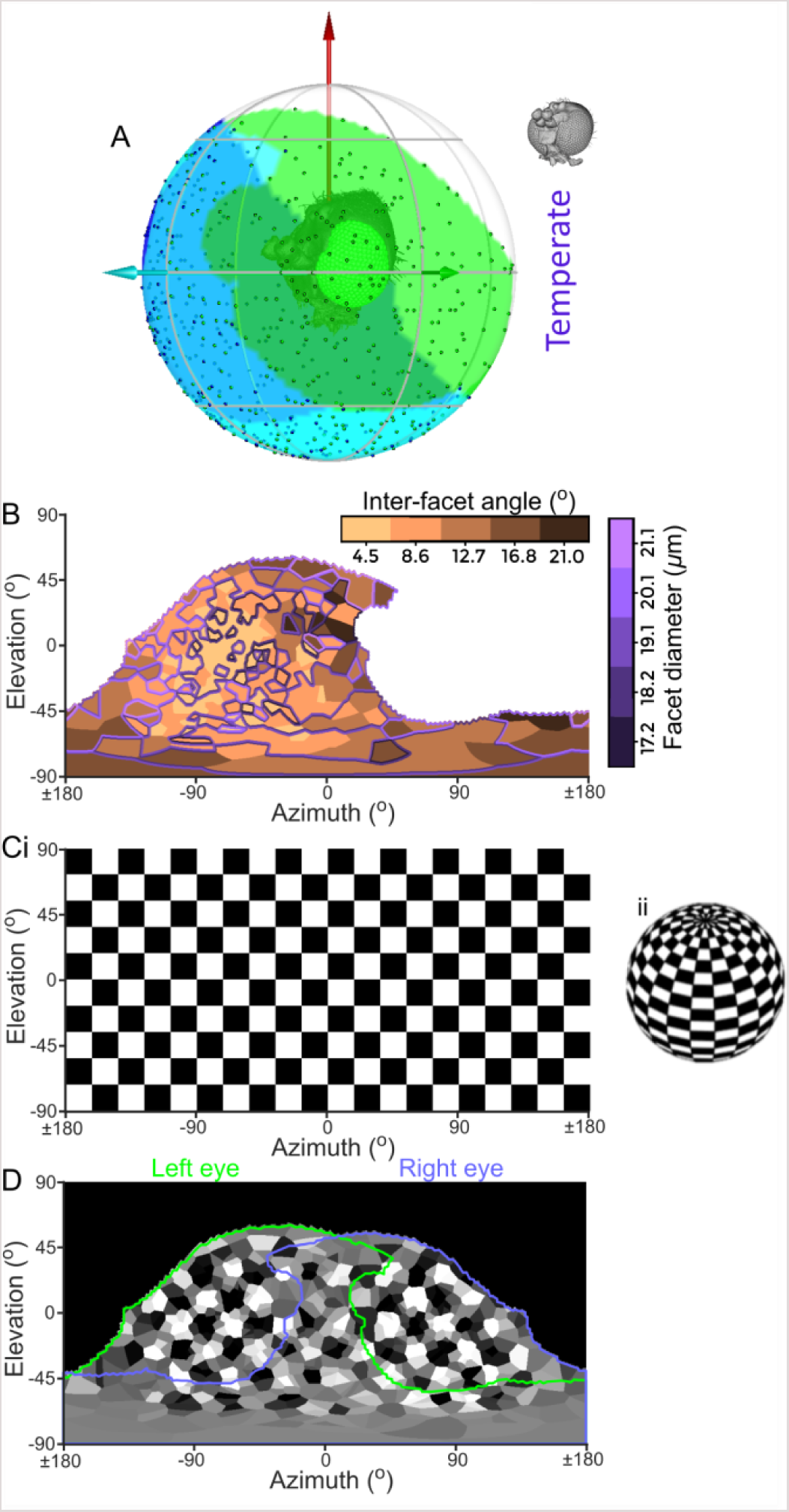
Examples of plotting options not used in the main text. (**A**) Head of a gnat plotted inside a sphere representing the world. Its left monocular, right monocular, and binocular CPs are represented by green, blue, and cyan shading on the sphere respectively. Points represent the intersection of the NVs of individual facets on the world sphere (as in Fig. 1Avii), and the green, cyan, and red arrows represent left lateral, frontal, and dorsal directions in the head-based reference frame. (**B**) Colour map of the IF angle projection on the world, overlaid with coloured contours representing the projected facet diameters. (**C**) Equirectangular projection (i) on a chequered sphere (ii) with 36° period. (**D**) visual simulation of the gnat vision as if it was located inside the chequered sphere. The green and mauve lines denote the limits of the CP for the left and right eyes respectively. The data used in this figure was from the temperate species *Neoplatyura modesta*.

